# Pre-TCR-Targeted Immunotherapy for T-cell Acute Lymphoblastic Leukemia

**DOI:** 10.1101/2024.05.29.596375

**Authors:** Patricia Fuentes, Marina García-Peydró, Marta Mosquera, Carmela Cela, Juan Alcain, Mireia Camós, Balbino Alarcón, María L. Toribio

## Abstract

Targeted immunotherapy for T-cell acute lymphoblastic leukemia (T-ALL), an aggressive tumor of developing T-cell progenitors, is an urgent unmet need, especially for relapsed/refractory (r/r) disease. Selective T-ALL targeting is challenging due to the shared antigen expression between leukemic and normal T cells. Here we identify the pre-TCR, a surface receptor essential for T-cell development, as a biomarker of leukemia-initiating cells (LICs) in human T-ALL. Loss-of-function genetic approaches demonstrate that pre-TCR signaling is necessary for LIC activity and tumor progression in pre-TCR^+^ T-ALL patient’s xenografts. Furthermore, we demonstrate the specific therapeutic targeting of pre-TCR with a monoclonal antibody against the invariant pTα subunit of the human pre-TCR, and validate an anti-pTα antibody-drug conjugate treatment as a potent immunotherapy for inhibiting LIC activity and tumor progression of T-ALL *in vivo*. These findings reveal the suitability of pre-TCR targeting as a promising therapy for the treatment of (r/r) patients with T-ALL expressing the pre-TCR.

## INTRODUCTION

T-cell acute lymphoblastic leukemia (T-ALL) is an aggressive neoplasia of developing T cells (1, 2), with significant rates of chemotherapy resistance and relapses, associated to dismal outcomes (20%-25% of pediatric and over 40% of adult patients) (3–5). Therefore, the development of novel therapies that selectively target leukemia initiating cells (LICs) of T-ALL, which are the ultimate responsible of relapses (6), is an urgent medical need. In recent years, the emergence of antibodies, antibody-drug conjugates (ADCs) and chimeric antigen receptor (CAR)-T cells, as highly potent antigen-targeted immunotherapies have revealed an unprecedented success for treatment of relapsed/refractory (r/r) B-cell malignancies like B-ALL (7–9). However, extension of this progress to T-ALL remains challenging, mainly because sharing of target antigens between normal and neoplastic T cells precludes tumor specificity, leading to aplasia of normal T-cells and increased risk of life-threatening immunodeficiency in treated patients (10–12). Identifying specific therapeutic targets involved in T-ALL pathogenesis and relapse, but not expressed in healthy T cells, is thus a crucial yet unmet goal.

T-ALL arises from the oncogenic transformation and developmental arrest of T-cell precursors at early stages of intrathymic maturation. This is a multistep process involving aberrant expression of oncogenic transcription factors and accumulating genetic alterations in key oncogenic, tumor suppressor and T-cell developmental pathways, which lead to uncontrolled cell survival and proliferation capacity (13). T-ALL can be classified into different subtypes based on the activation of specific T-ALL oncogenes and associated gene-expression profiles that correlate with the stage of arrest in T-cell development (14).

NOTCH1 is a critical developmental pathway involved in T-cell lineage commitment, expansion and differentiation (15,16). Somatic activating mutations in NOTCH1 have been identified in a high frequency of T-ALL cases (>60%) from all major molecular subtypes, resulting in overexpression of constitutively active NOTCH1 (ICN1) (17). Comprehensive studies on the genomic landscape of T-ALL in patients confirmed the dominant presence of NOTCH1 as a driver mutation (13,18). In fact, NOTCH1 activation is an early hallmark of T-cell leukemogenesis and a key regulator of LIC activity in different T-ALL subtypes (19–21), suggesting that deregulated transcription of NOTCH1 target genes and aberrant activation of downstream proliferation pathways critically contribute to NOTCH1-dependent T-ALL pathogenesis (13,18,22). Given the high prevalence of NOTCH1 mutations in T-ALL, and the dependence of T-ALL cases on NOTCH1 function, understanding the role of molecular effectors downstream of NOTCH1 in T-ALL should provide key information for developing specific targeted therapies for this disease.

One of the NOTCH1 targets with a prominent role in T-cell development is the pre-T cell receptor (pre-TCR). NOTCH1 transcriptionally regulates *PTCRA* (23, 24), the gene encoding the invariant pTα chain that associates with the TCRβ chain and forms the pre-TCR in developing T cells (25–27). In both mice and humans, the pre-TCR is expressed transiently in intrathymic T-cell precursors, and identifies a critical developmental checkpoint, known as β-selection, which leads to the survival and expansion of pre-T cells with functional TCRβ gene rearrangements, and finally results in their differentiation into mature T cells expressing the TCRαβ (27–30). The prominent expansion function of the pre-TCR during T-cell development suggested its participation in leukemogenesis. However, genetic approaches on animal models have generated variable results, indicating that pre-TCR function can be either important (31–34) or dispensable (35, 36) for T-ALL pathogenesis depending of the particular driver oncogene model used. More importantly, very little is known about pre-TCR expression in human T-ALL. Seminal studies have proposed that half of the TCRαβ-lineage T-ALL cases in patients might express a pre-TCR (37), but the significance of pre-TCR signaling in human T-ALL is unknown, and the involvement of pre-TCR in the maintenance and progression of T-ALL in patients remains largely undefined. Herein, we addressed such important questions based on an *in vivo* model of human NOTCH1-induced T-ALL pathogenesis (38), in combination with pre-TCR loss-of-function approaches in T-ALL patient-derived xenografts (PDX), and preclinical assays of immunotherapy based on administration of a monoclonal antibody (mAb) specific for the extracellular domain of the pTα chain of the human pre-TCR (39). Our study provides proof-of-concept data for the relevance of pre-TCR signaling in human T-ALL leukemia-initiating cell (LIC) activity and tumor progression, uncovering the suitability of the pre-TCR as a promising therapeutic target for treatment of (r/r)T-ALL patients.

## RESULTS

### Human T-ALL pathogenesis induced by active NOTCH1 involves a developmental arrest at the β-selection checkpoint

To study the multistep pathogenic process underlying T-ALL generation in patients, we recently extended the reported model of NOTCH1-induced murine T-ALL (40) to humans (38). To this end, immunodeficient NSG mice were intravenously (i.v.) injected with human cord blood (CB) CD34^+^ hematopoietic progenitor cells (HPCs) transduced with constitutively active NOTCH1 (ICN1) and GFP as cell tracer (**Figure 1A**). Injected mice generated *de novo* a clonal human T-ALL that recapitulated T-ALL in patients, comprising a major (>95%) population of CD4+ CD8+ double positive (DP) cells with variable expression levels of surface CD3 (38 and **Figure 1B**). Detailed flow cytometry revealed that a significant fraction of DP leukemic cells in all animals (up to 60% at 20-40 weeks post-transplant) expressed low CD3 surface levels (CD3^lo^) and were negative surface αβ and γδ TCR (TCR^neg^) (**Figure 1B**). This phenotype was compatible with a developmental arrest at the β-selection DP intrathymic stage where the pre-TCR is expressed (26). Next, we sought to assess the pathogenic potential of the arrested pre-T-like cells when transferred to recipient mice. We found that numbers of DP CD3^lo^ TCR^neg^ cells increased progressively in first and secondary recipients, where they were able to serially transfer an aggressive T-ALL (**Figure 1C, D**), indicating that they display a selective *in vivo* growth advantage characteristic of leukemic cells with LIC potential. Therefore, we concluded that NOTCH1-induced human T-ALL cells arrested at the DP CD3^lo^ TCR^neg^ β-selection stage may comprise T-ALL LIC cells.

**Figure 1.**
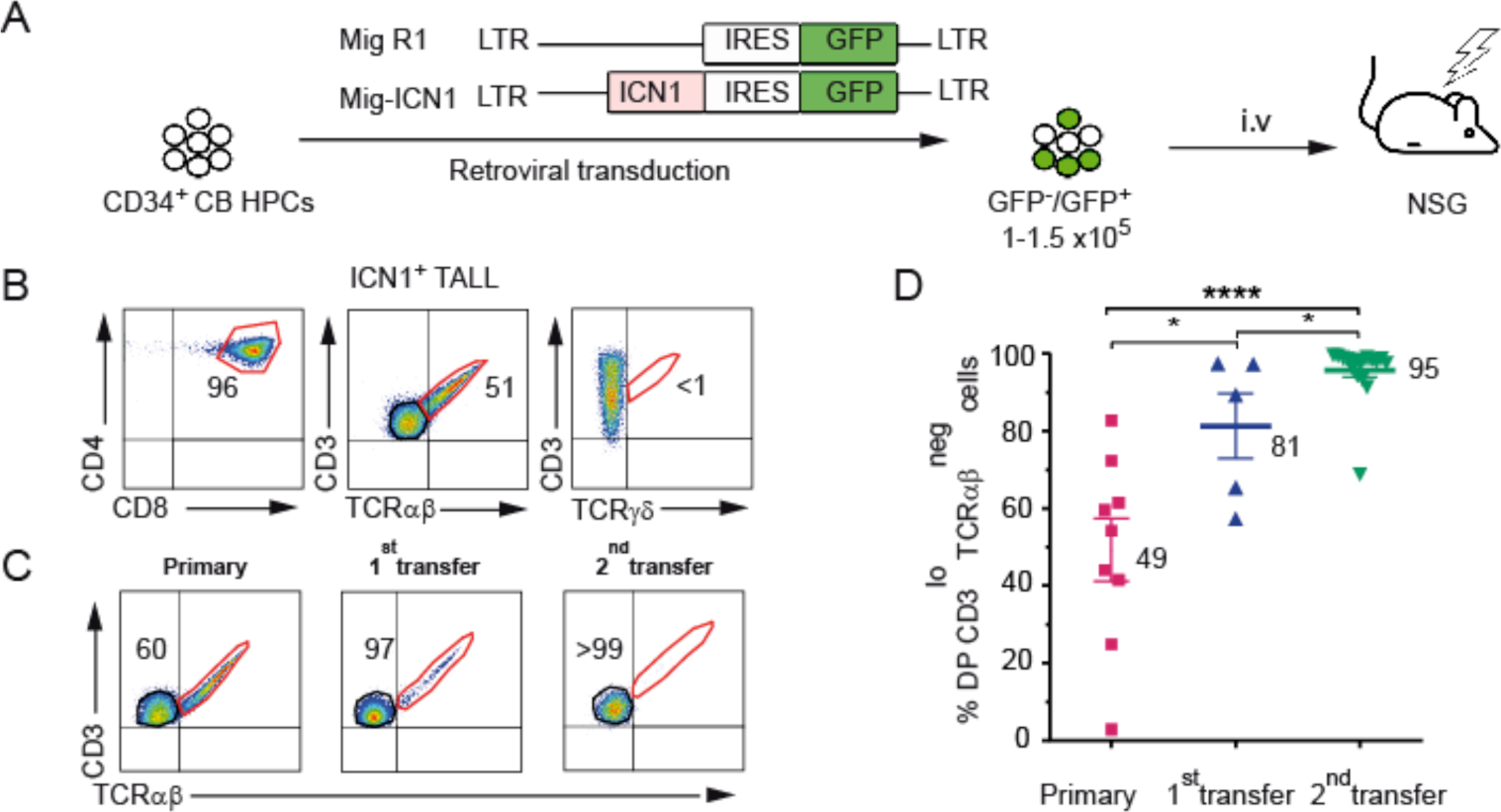
Human T-ALL pathogenesis induced by active NOTCH1 involves a developmental arrest at the β-selection checkpoint. (A) Schematic representation of experimental design for *de novo* generation of human T-ALL. Sublethally irradiated NSG mice were subjected to i.v. injection with human CB CD34^+^ HPCs (1-1.5×10^5^ cells per mice) retrovirally transduced with either active NOTCH1 (ICN1) and GFP, or GFP alone as control. (B) Representative flow cytometry expression of CD4, CD8, CD3, TCRαβ and TCRγδ by ICN1^+^ human T-ALL cells infiltrating the BM of diseased mice in (A) at 27-weeks post-transplant. (C) Representative CD3 vs TCRαβ expression of human ICN1^+^ T-ALL cells recovered from diseased mice in (A) (primary), or from NSG mice after serial transplantation with BM from primary mice (1^st^ transfer and 2^nd^ transfer). Results correspond to T-ALL cells isolated from the BM at 39, 8 and 9 weeks post-transplant, respectively. (D) Mean percentages ± SEM of human CD8^+^CD4^+^ DP, CD3^low^ TCRαβ^neg^ ICN1^+^ cells infiltrating the BM of either primary diseased mice, or serially transplanted mice (9 mice per group from 2 independent experiments) as in (C). \**P* < 0.05; \*\*\*\**P* < 0.0001.

### Patient T-ALL cells arrested at the β-selection checkpoint display LIC potential

To evaluate the clinical relevance of the DP CD3^lo^ TCR^neg^ leukemic cells generated in the human T-ALL model, we examined whether an equivalent population could be identified in T-ALLs from patients. We analyzed the phenotype of a cohort of T-ALL human samples belonging to several biological subgroups representative of arrest at different developmental stages, as defined by the European Group for Immunological Characterization of Leukemias (EGIL) (14) (**Table S1**). According with pioneering studies by Macintyre’s group in a large T-ALL cohort (37), about 50% of T-ALL samples (9 out of 19) in our cohort included variable proportions of cells with a CD3^lo^ TCR^neg^ phenotype, which lacked surface αβ and γδ TCR, but displayed cytoplasmic TCRβ (cTCRβ) (**Figure 2A, Table S1**), and may thus express the pre-TCR (26, 37). Most samples showed a cortical T-ALL phenotype, while some belonged to the pre-T biological subtype, and all of those analyzed for NOTCH1 activation, exhibited intracellular active ICN1 (**Table S1**). Therefore, half of human T-ALL cases include a CD3^lo^ TCR^neg^ population arrested at the β-selection stage, which seems equivalent to the LIC population characterized in the ICN1-induced human T-ALL model (**Figure 1A**). Accordingly, they showed to display a selective *in vivo* growth advantage, given that, regardless of the initial proportion of CD3^lo^ TCR^neg^ cells in the patient sample, their relative numbers increased progressively in the BM and spleen of serially transplanted NSG mice (**Figure 2B, C**). Confirming the LIC function of CD3^lo^ TCR^neg^ cells, limiting dilution PDX assays showed that LIC activity was significantly enriched *in vivo* (40-fold) in FACS-sorted CD3^lo^ TCRαβ^neg^ cells, compared with CD3^+^ TCRαβ^+^ cells isolated from the very same T-ALL patient sample (**Figure 2D, E**), and LIC frequencies inversely correlated with overall survival of transplanted mice (**Figure 2F**). Together, these results indicate that half of human T-ALL cases may include variable proportions of leukemic cells with LIC potential, which display the phenotype of thymocytes at the pre-TCR^+^ β-selection stage.

**Figure 2.**
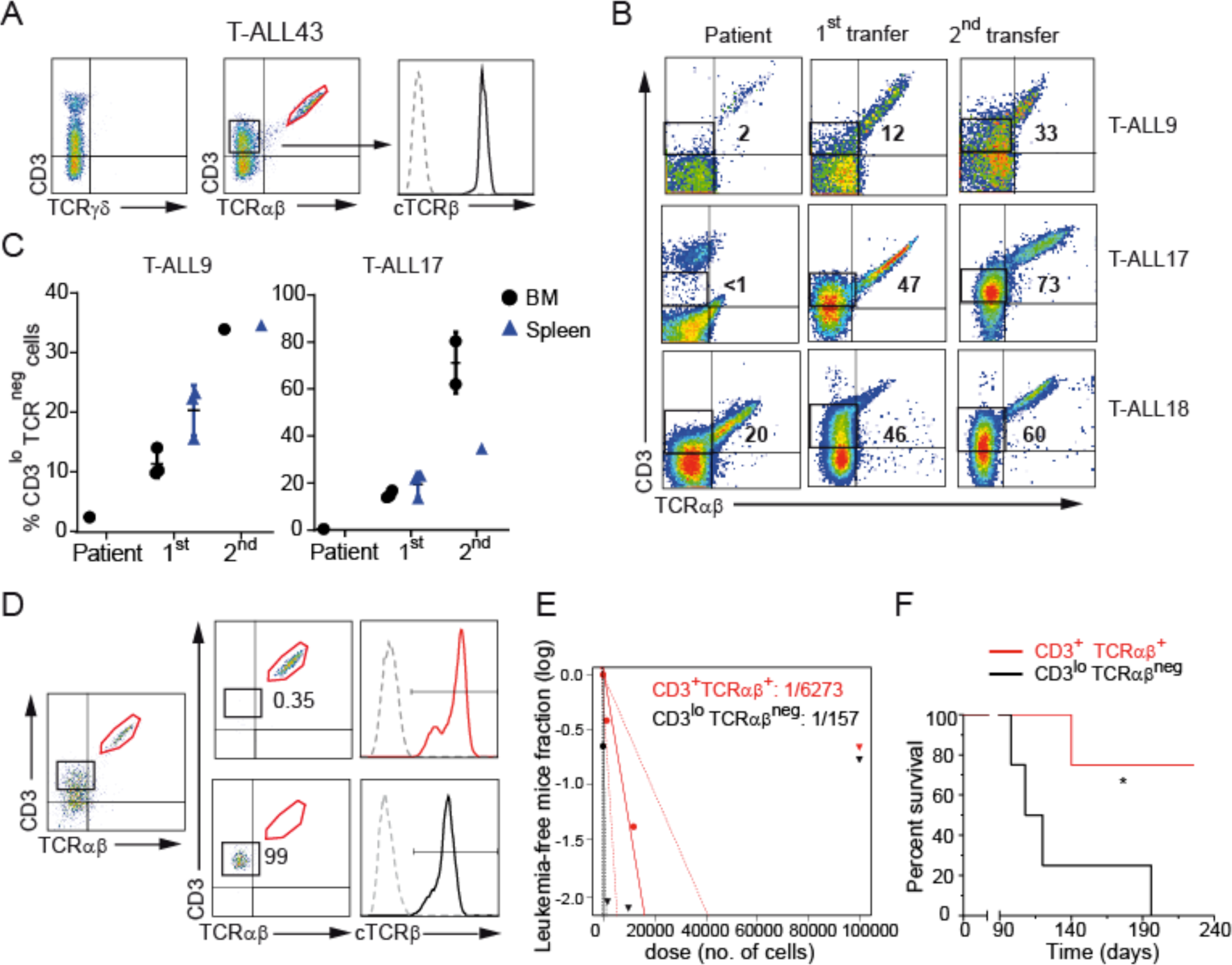
T-ALL patient cells arrested at the β-selection checkpoint display LIC potential. (A) CD3, TCRαβ and TCRγδ expression of a T-ALL patient sample (T-ALL43) representative of developmental arrest at the β-selection checkpoint. Expression of cytoplasmic TCRβ (cTCRβ) by gated CD3^lo^ TCR^neg^ cells (middle histogram) is shown in the monoparametric histogram on the right. Dashed histogram shows background staining with an irrelevant isotype-matched antibody. (B) CD3 vs TCRαβ expression displayed by primary PB T-ALL cells from 3 patients (T-ALL9, T-ALL17, T-ALL18) analyzed by flow cytometry at the time of diagnosis (Patient), or after serial transplantation into consecutive NSG mice (1^st^ and 2^nd^ transfer). Analyses of cells engrafting the BM of transplanted mice were performed at 7- to 22-weeks post-transplant. Numbers indicate percentages of gated CD3^low^ TCRαβ^neg^ cells. (C) Mean percentages ± SEM of human CD3^lo^ TCRαβ^neg^ cells from T-ALL9 and T-ALL17 patient samples engrafting the BM and spleen of serially transplanted mice in (B) (1-3 mice per group). Percentages at the time of diagnosis (Patient) are shown for comparison. (D) Sorting strategy used to isolate CD3^lo^ TCRαβ^neg^ and CD3^+^ TCRαβ^+^ T-ALL17 patient cells recovered from spleen xenografts of NSG mice at 3.5-weeks post-transplant (left histograms). CD3 *vs* TCRαβ and cTCRβ expression of sorted populations is shown in the middle and right histograms, respectively. (E) LIC potential of CD3^+^ TCRαβ^+^ (red) and CD3^lo^ TCRαβ^neg^ (black) human T-ALL17 sorted populations shown in (D) and transplanted into NSG mice under limiting dilution conditions (from 10^5^ to 10 cells per mouse, 3-4 mice per dose). LIC frequency was calculated using ELDA software by recording numbers of leukemia-free mice in the PB by 32 weeks post-transplant. (F) Kaplan-Meier survival curves of mice transplanted in (E) with 10^3^ FACS-sorted CD3^+^ TCRαβ^+^ (red) or CD3^lo^ TCRαβ^neg^ (black) human T-ALL17 cells. \**P* <0.05.

### Functional pre-TCR expression is a biomarker of human T-ALL cells with LIC potential

Next, we ought to confirm that CD3^lo^ TCR^neg^ cells with LIC function do in fact express the pre-TCR at the cell surface. To this end, we took advantage of a mAb (K5G3) generated against the pTα component of the human pre-TCR (39), that we previously characterized by its reactivity with JR.pTα, a TCRα-deficient Jurkat cell line (JR3.11) (41) stably transfected with human pTα. Flow cytometry analysis showed that the anti-pTα mAb was reactive with CD3^lo^ TCR^neg^ SupT1 human leukemic pre-TCR^+^ control cells (39), but not with CD3^+^ HPB-ALL and Jurkat T-ALL cells expressing the conventional TCRαβ (**Figure 3A**). In addition, we consistently observed a specific, although very low reactivity of the anti-pTα mAb with different CD3^lo^ TCR^neg^ T-ALL patient samples, which also displayed low CD3 surface levels, significantly lower than those found on TCRαβ^+^ cells (**Figure 3B**). Low anti-pTα mAb reactivity was observed as well in DP CD3^lo^ TCR^neg^ cells from a 17-weeks-old human fetal thymus sample, and in cells developing *in vitro* either from human CB HPCs co-cultured with OP9-Jag2 stroma, or from human early thymic progenitors (ETPs) cultured in artificial thymic organoids (ATOs) (**Figure 3C**). Collectively, our data confirm that the pre-TCR is expressed at low levels on CD3^lo^ TCR^neg^ patient T-ALL cells arrested at the β-selection stage, as previously shown in their physiological intrathymic counterparts (27,42).

**Figure 3.**
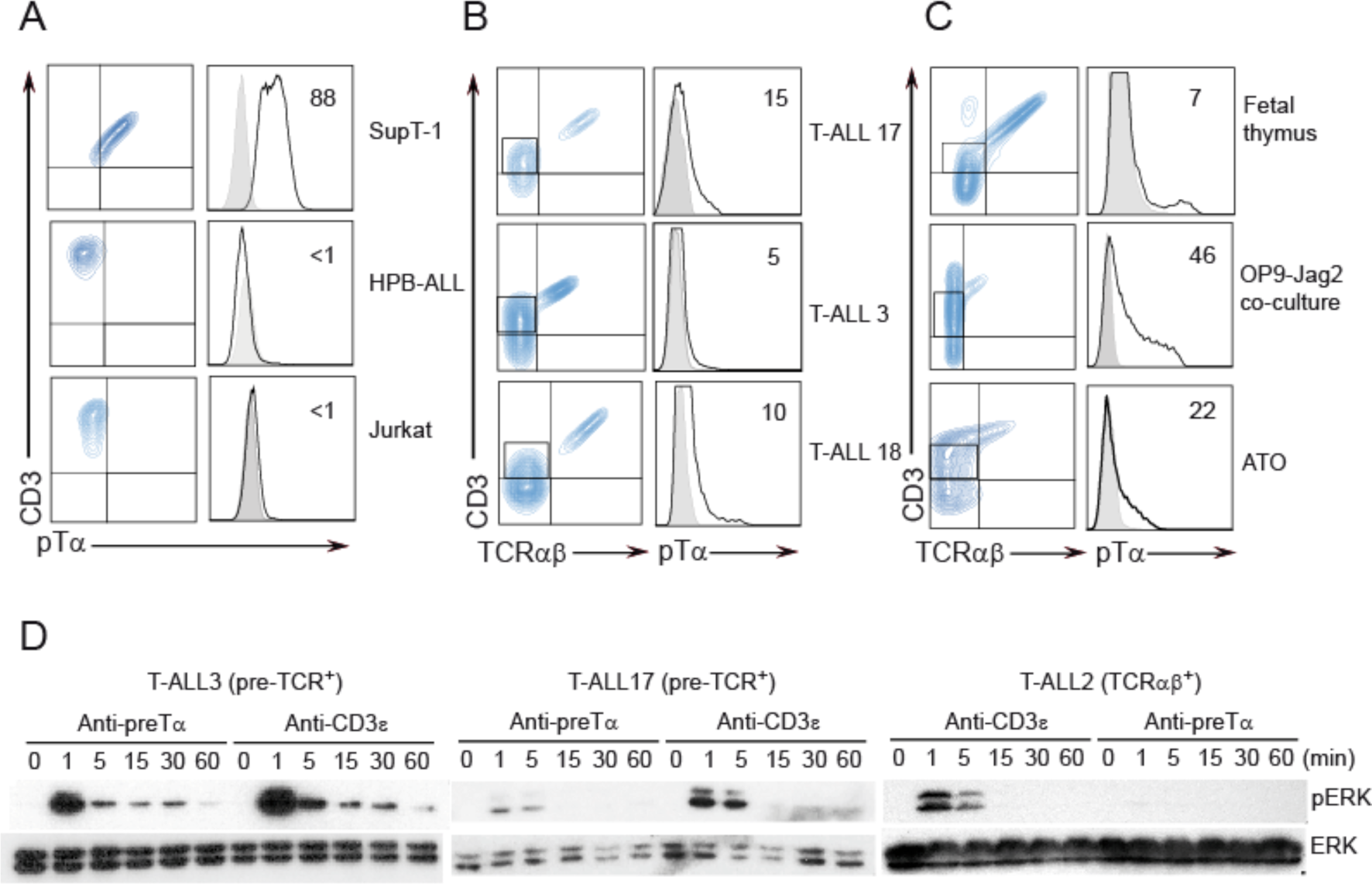
Expression of low but functional levels of surface pre-TCR in patient T-ALL. (A) Flow cytometry analysis of CD3 *vs* pre-TCR expression in pre-TCR-expressing SupT1, and TCRαβ-expressing HPB-ALL and Jurkat T-ALL cell lines, as assessed by reactivity with anti-CD3 and anti-pTα mAbs (left biparametric histograms). Monoparametric histograms on the right show surface pre-TCR expression levels measured with an anti-pTα mAb. Numbers indicate percentages of positive cells. Shaded histograms show background staining with an irrelevant isotype-matched antibody (B) Phenotype of 3 patient T-ALL samples (T-ALL3, T-ALL17, T-ALL18) showing CD3 *vs* TCRαβ expression (left) and pre-TCR expression levels of electronically gated CD3^lo^ TCRαβ^neg^ T-ALL cells analyzed with an anti-pTα mAb (right). (C) Representative CD3 *vs* TCRαβ expression of either primary human thymocytes from a 17-weeks-old fetal thymus sample (upper left), or developing T cells derived either from human CB HPCs after 70 days of culture onto OP9-Jag2 cells (middle left), or from CD34^+^ human ETPs after 18 days of culture in ATOs (bottom left). Pre-TCR expression levels of electronically gated CD3^lo^ TCRαβ^neg^ cells analyzed with an anti-pTα mAb are shown in the monoparametric histograms on the right. Numbers indicate percentages of positive cells relative to background staining with an isotype-matched mAb. (D) Immunoblot analysis of ERK activation in pre-TCR^+^ (T-ALL3 and T-ALL17) or TCRαβ^+^ (T-ALL2) patient T-ALLs, stimulated either with anti-pTα or with anti-CD3ε mAbs for the indicated times. Total ERK expression is shown as loading control.

A key question is whether pre-TCR expression levels that are barely detectable by flow cytometry, are functionally competent in patient leukemias, as occurs in developing thymocytes. To investigate this issue, pre-TCR activation was examined by means of ERK signaling induction (43) in western blotting of primary T-ALLs (T-ALL3 and T-ALL17), following *in vitro* pre-TCR activation with the anti-pTα or with an anti-CD3ε mAb. A TCRαβ^+^ T-ALL sample (T-ALL2), included in the study for comparison, showed ERK activation only in response to anti-CD3ε, confirming the specificity of the anti-pTα mAb (**Figure 3D**). In contrast, both T-ALL3 and T-ALL17 showed similar ERK phosphorylation kinetics in response to either anti-pTα or anti-CD3ε mAbs (**Figure 3D**), indicating that the low levels of pre-TCR expressed on the surface of CD3^lo^ TCR^neg^ T-ALL patient cells are functionally competent. Thus, we concluded that surface expression of a functional pre-TCR is a biomarker of LIC cells in ∼50% of T-ALL patients.

### Pre-TCR signaling is required for LIC activity and tumor progression of patient T-ALL

The expression of a functional pre-TCR on human T-ALL cells with LIC potential raises the possibility that pre-TCR signaling participates in the proliferation and function of human T-ALL LICs. To directly investigate this issue, we analyzed the LIC potential and tumor progression of patient T-ALL cells in which pre-TCR function was abrogated by gene-silencing of a critical pre-TCR functional adaptor, CMS, which binds to the human pTα cytoplasmic domain and is required for pre-TCR activation (44). Silencing of the CMS-encoding *CD2AP* gene was first confirmed by western blotting of CMS expression in the pre-TCR^+^ SupT1 cell line upon transduction with several short hairpin (sh) RNAs (shCMS), as compared with non-transduced cells (**Figure S1A**). Then, the best-performing shCMS (shCMS4), or a control scrambled shRNA (shSC), were transduced into the pre-TCR^+^ JR.pTα cell line and PLCγ1 activation, which is crucial for pre-TCR signal transduction (45,46), was analyzed upon anti-CD3-induced receptor crosslinking as readout of pre-TCR function. These assays confirmed that CMS silencing resulted in defective pre-TCR signaling, as indicated by impaired PLCγ1 activation and NFAT activity, when compared with controls (**Figure S1B, C**). More importantly, CMS silencing also led to deficient pre-TCR signaling *in vivo*, as illustrated by the impaired generation of human thymocytes in Rag2^-/-^xγc^-/-^ mice at week 5 post-transplantation with human ETPs transduced with shCMS, as compared with control mice transplanted with shSC-transduced ETPs (**Figure S1D, 1E, F)**. Of note, no significant differences in thymus reconstitution were observed in both groups of mice before week 3 post-transplant, when most progenitors are reaching the DP CD3^lo^ β-selection stage **(Figure S1E, 1F-left)**; but only control shSC ETPs developed efficiently into β−selected DP CD3^+^TCRαβ^+^ cells by 5 weeks (**Figure S1E, 1F-right**). Therefore, impaired pre-TCR signaling resulting from CMS silencing specifically hampers pre-TCR-mediated transition through the β-selection checkpoint, this compromising the development of downstream TCRαβ+ cells.

Having validated the CMS silencing approach for pre-TCR function inhibition, we next evaluated the *in vivo* LIC potential displayed by T-ALL patient samples (T-ALL5 and T-ALL9) with a compromised pre-TCR function (**Figure 4A).** To this end, we analyzed tumor progression in NSG mice i.v. injected with T-ALL cells transduced with either shCMS plus GFP or shSC plus GFP as control **(Figure 4B, D**). Control mice showed significant leukemic engraftment and progression, with clear signs of disease by weeks 11 and 13 post-transplantation with T-ALL9 and T-ALL5, respectively. At this time, mice showed leukemic infiltration of distinct organs including BM, thymus, brain, liver and spleen. In contrast, expansion of shCMS-transduced T-ALL cells was significantly impaired in all peripheral organs of transplanted mice, according with an impaired LIC potential (**Figure 4C, D, E**). Supporting the specificity of shCMS on pre-TCR^+^ T-ALL cells, LIC potential of patient B-ALL cells (B-ALL9) transduced with shCMS remained intact, as leukemic cells were capable of expanding and progressing as efficiently as shSC-transduced B-ALL controls in transplanted mice, and no significant differences were found between both groups in either the PB or the BM, spleen and liver, at 4.5- and 7-weeks post-transplant, respectively (**Figure 4F-H**). Collectively, these results provide evidence that defective pre-TCR signaling selectively impairs cell proliferation and tumor progression of primary human pre-TCR^+^ T-ALLs *in vivo*, highlighting a crucial role for pre-TCR function in LIC activity of human T-ALL. In addition, our data raise the possibility that the pre-TCR may also have an active role in the pathogenesis of pre-TCR^+^ T-ALLs. Supporting this view, we found that the ICN1 construct shown to generate human pre-TCR^+^ leukemias (**Figure 1**) was unable to induce murine T-ALL when BM HPCs from CD3ε C80G mutant mice were ICN1-transduced and inoculated into NSG recipient mice (**Figure S2**). The C80G mutation in CD3ε has been shown to cause a total arrest of normal thymocyte differentiation at the pre-TCR stage due to a blockade of pre-TCR signaling (47). In contrast to C80G mutant HPCs, WT BM HPCs transduced with the ICN1 construct developed an aggressive leukemia in NSG mice as early as at 3 weeks post-transplant, leading to peripheral organ infiltration and a marked splenomegaly, and resulting in a median survival of 5-weeks (**Figure S2**). Therefore, the implication of the pre-TCR in T-ALL LIC activity could be extended to T-ALL pathogenesis in human T-ALL cases arrested at the pre-TCR^+^ β-selection stage. Collectively, these data point to the pre-TCR as a promising therapeutic target for T-ALL.

**Figure 4.**
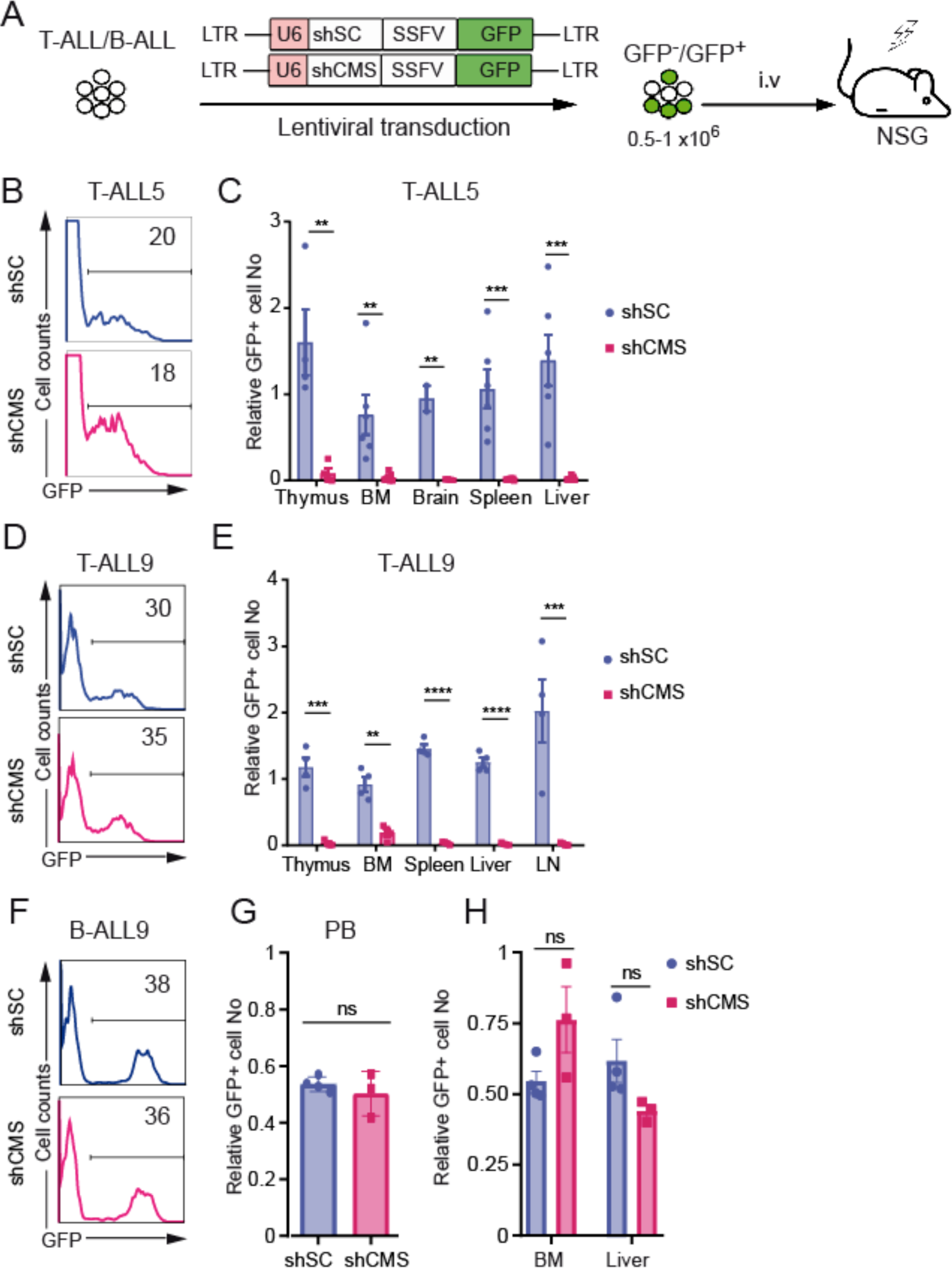
Pre-TCR signaling is required for LIC activity and tumor progression of human T-ALL. (A) Schematic diagram of experimental design for abrogation of pre-TCR function in patient T-ALL cells. Patient T-ALL (T-ALL5 and T-ALL9) or B-ALL (B-ALL401) cells recovered from BM xenografts of NSG mice were transduced with a lentiviral vector encoding a shRNA against CMS (shCMS) and GFP as cell tracer, or a scramble RNA (shSC) and GFP, as control, prior to transplantation into sublethally irradiated NSG mice. (B, D, F) Transduction efficiencies of transplanted T-ALL5 (B), T-ALL9 (D) and B-ALL9 (F) were analyzed by flow cytometry. Numbers indicate percentages of transduced (GFP^+^) cells. (C, E, G, H) Relative numbers of T-ALL5 (C), T-ALL9 (E) and B-ALL9 (G, H) cells transduced with shCMS or shSC, engrafting the indicated organs of NSG mice euthanized when they presented advanced symptoms of disease (13, 11 and 7 weeks post-transplant for T-ALL5, T-ALL9 and B-ALL9, respectively). Data are shown as mean ± SEM percentages of CD45^+^GFP^+^ engrafted cells in 4 to 8 mice per group, normalized to percentages of GFP^+^ injected cells (B, D and F). \*\**P* < 0.01; \*\*\**P* < 0.001; \*\*\*\**P* < 0.0001.

### Preclinical validation of pre-TCR therapeutic targeting for T-ALL treatment

To directly evaluate the T-ALL therapeutic potential of pre-TCR targeting, we used an *in vivo* preclinical assay consisting on the administration of our anti-pTα mAb in a PDX model. Patient T-ALL (T-ALL-3) cells were i.v. injected into sublethally irradiated (1.5 Gy) SCID mice, which were subjected to intraperitoneal (i.p.) administration of either the anti-pTα mAb or an isotype-matched (IgG2a) control (10mg/Kg, twice a week) during 10 weeks, starting at week 3 post-transplant, when the disease was clearly established (>1% of total PB white cells) (**Figure 5A**). Sequential analysis of PB revealed that T-ALL engraftment increased dramatically to 90% during the first 8 weeks post-transplant in control animals, while only 5% of leukemic cells engrafted the PB of anti-pTα-treated mice at this time (**Figure 5B**). Thereafter, T-ALL progression was significantly delayed in the PB of mice treated with the anti-pTα mAb, as compared with control mice, which showed clear symptoms of disease by 11 weeks post-transplant and should be euthanized. Accordingly, the median survival of anti-pTα-treated mice was significantly prolonged (>50%) when compared with that of isotype control-treated mice (**Figure 5C**), this validating the therapeutic potential of the anti-pre-TCR targeting strategy based on anti-pTα mAb administration.

**Figure 5.**
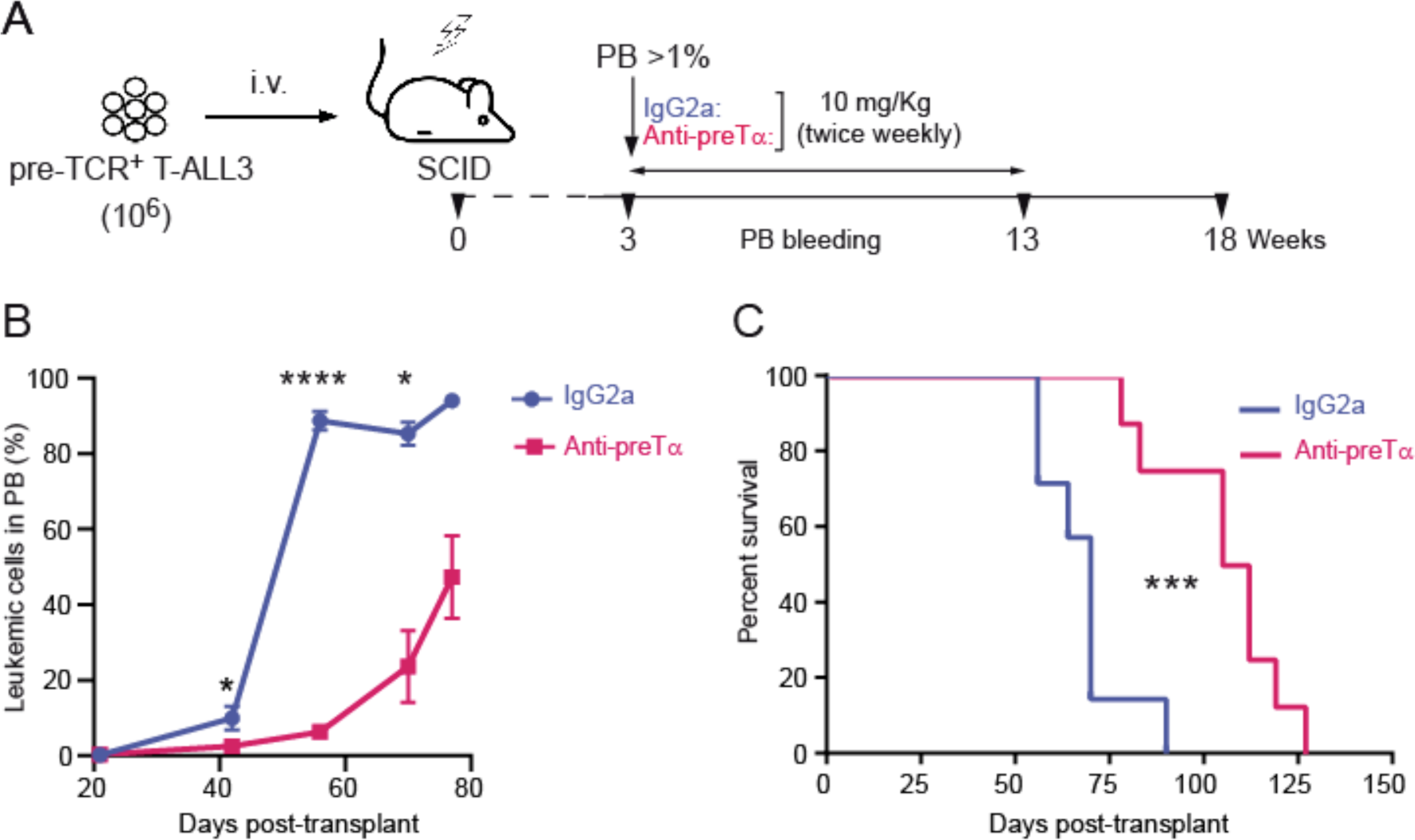
Preclinical validation of pre-TCR therapeutic targeting for T-ALL treatment. (A) Schematic design of *in vivo* experimental treatment of patient T-ALL with an anti-pTα mAb. Sublethally irradiated SCID mice subjected to i.v. iinjection with T-ALL3 patient cells (10^6^ cells per mice) were serially i.p. injected (twice weekly for 10 weeks) with either anti-pTα mAb or isotype-matched control (IgG2a, 10mg/Kg), starting at week 3 post-transplant, when T-ALL engraftment in PB was >1%. B) Percentages of T-ALL3 cells infiltrating the PB of mice at the indicated days post-transplant, upon treatment with anti-pTα mAb or IgG2a control. Data are shown as mean percentages ± SEM from 6 (control-) or 8 (anti-pTα-) treated mice. (C) Kaplan-Meier survival curves of mice in (B). \**P* <0.05; \*\*\**P* < 0.001; \*\*\*\**P* < 0.0001.

### Targeted immunotherapy with anti-pTα antibody drug conjugates efficiently impairs tumor progression of human pre-TCR+ T-ALL *in vivo*

The impaired progression, though not complete elimination, of T-ALL in anti-pTα mAb-treated animals prompted us to search for strategies to maximize the efficacy of this targeted immunotherapy. ADCs represent a compelling clinical option particularly efficient for targeting surface receptors endowed with dynamic internalization properties (48), like those exhibited by the pre-TCR (42,44,49). In fact, we previously reported that the anti-pTα mAb is internalized after binding to the pre-TCR and traffics into lysosomes (38), indicating that it could be optimal for delivery of cytotoxic drugs. Thus, the anti-pTα mAb was subjected to drug conjugation using either non-cleavable or cleavable linkers and either Mertansine (DM1) or Monomethyl auristatin E (MMAE), respectively (**Figure 6A**), two toxins which disrupt the cytosolic microtubule network leading to blocked mitosis, impaired proliferation and apoptosis (48). The resulting anti-pTα ADCs were functionally validated *in vitro* for their pre-TCR binding capacity, as they bound to the CD3 complex on SupT1 cells and on JR.pTα cell lines generated from TCRα^-^ JR3.11 cells transduced with a pTα-GFP MigR1.1 retroviral vector (44). In contrast, both ADCs failed to react with TCRαβ^+^ Jurkat cells and with JR3.11 cells transduced with a GFP retroviral control (JR.MigR1), as observed with the conventional anti-pTα drug-free mAb (**Figure 6B**). Flow cytometry also confirmed that both anti-pTα ADC variants displayed optimal features for cargo intracellular delivery, as they induced a rapid internalization of surface pre-TCR (>60-70% in 5 min), with kinetics similar to those observed with the uncoupled anti-pTα mAb (**Figure 6C**). Moreover, both ADCs impaired T-ALL cell cycle progression *in vitro* by inducing G2/M cell cycle arrest of pre-TCR+ SupT1 and JR.pTα cells (**Figure 6D, E**), which resulted in a significant and specific decrease of cell proliferation (**Figure 6F),** but they had no effect on pre-TCR^-^ HPB-ALL and JR.MigR1 cells.

**Figure 6.**
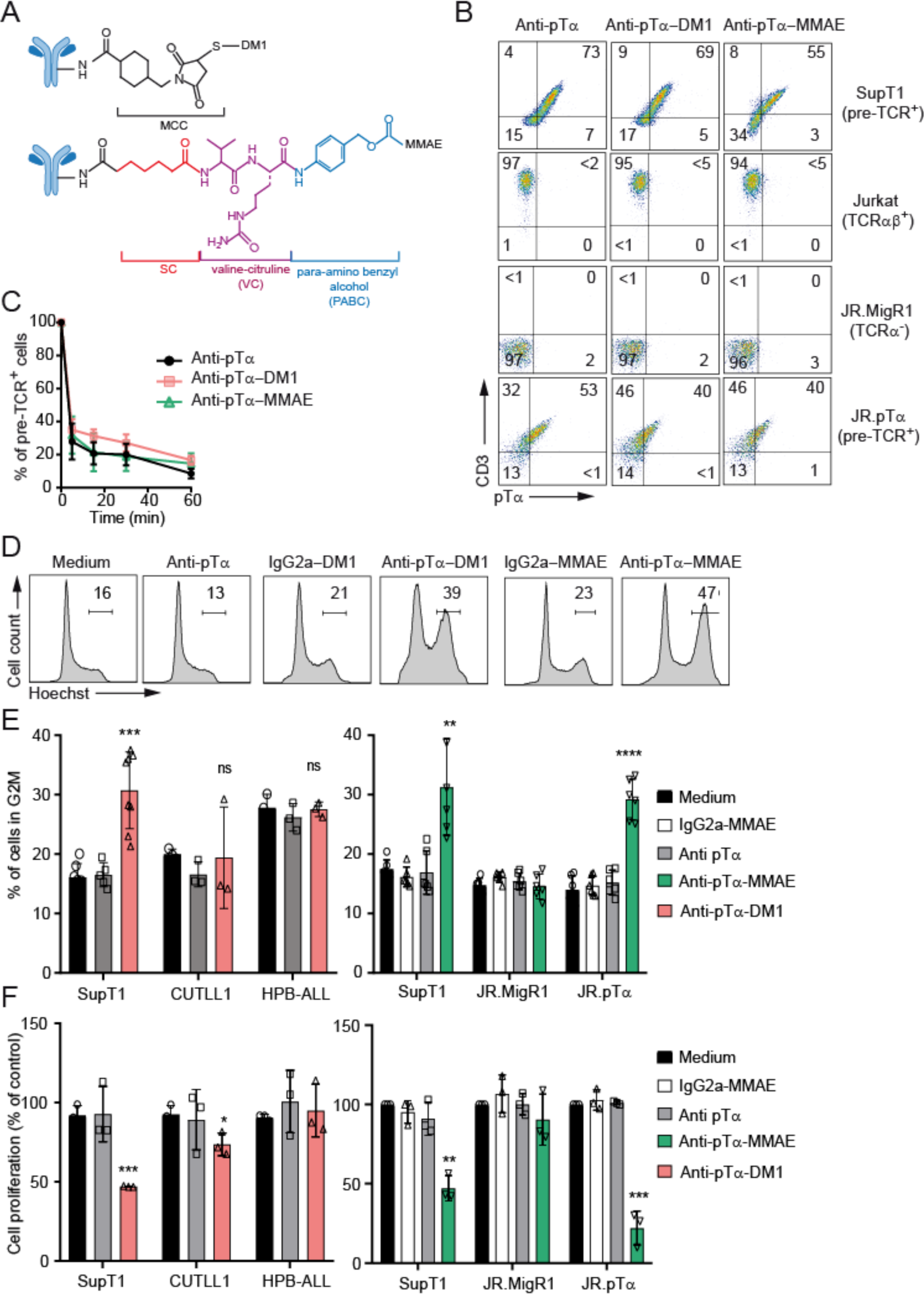
Anti-pTα ADCs induce pre-TCR internalization and impair cell cycle progression and cell proliferation of pre-TCR^+^ T-ALL cells. (A) Illustration of the modular nature of anti-pTα-DM1 and anti-pTα-MMAE ADCs, showing non-cleavable (MCC) and cleavable linkers (DC-VC-PABC) used. (B) Surface pre-TCR expression levels detected on SupT1 and JR.pTα pre-TCR^+^ cell lines by flow cytometry, using anti-CD3 in combination with either drug-free anti-pTα mAb, or anti-pTα-DM1 or anti-pTα-MMAE ADCs. Pre-TCR-negative Jurkat (TCRαβ^+^) and mock-transduced JR3.11 (JRMigR1, TCRα^-^) cell lines were assayed as negative controls. (C) Pre-TCR internalization in SupT1 cells pulsed with drug-free anti-pTα mAb or anti-pTα ADCs at 4°C for 30min and incubated for the indicated times at 37°C. Results are shown as mean percentages ± SEM of pre-TCR^+^ cells detected by flow cytometry with a secondary anti-mouse IgG-FITC antibody normalized to time 0, (n=3). (D) Representative cell cycle profiles of SupT1 pre-TCR^+^ cells left untreated (medium) or treated with either drug-free anti-pTα mAb, anti-pTα-DM1 or anti-pTα-MMAE ADCs, or their respective isotype-matched controls for 4 days. Percentages of gated cells in the G2/M cell cycle phases are shown. (E) Mean percentages ± SEM of cells in the G2/M phases of the cell cycle within the indicated pre-TCR^+^ (SupT1, JR.pTα) and pre-TCR-negative (CUTLL1, HPB-ALL, JR3.MigR1) cell lines, treated as in (D), (n≥3). (F) Relative cell proliferation of the indicated leukemic cells recovered after in vitro treatment as in (D). Data are shown as mean ± SEM percentages of recovered treated cells relative to untreated cells, (n≥3). *P < 0.05, **P < 0.01, ***P < 0.001, ****P < 0.0001.

Next, we sought to investigate the *in vivo* efficacy of a specific immunotherapy based on anti-pTα ADC administration to NSG mice xenografted with patient pre-TCR^+^ T-ALLs (T-ALL3 or T-ALL42). Treatment consisting of i.p. injection twice a week for 3 weeks of the anti-pTα-DM1 ADC, was initiated when the disease was clearly established in mice receiving T-ALL3 (>1% T-ALL3 cells in PB; 8 weeks post-transplant). Alternatively, 3 cycles of established induction chemotherapy based on vincristine, dexamethasone and L-asparaginase (VxL) (50) were administered as control (**Figure 7A**). PB examination of vehicle-treated (PBS) control mice revealed a massive expansion (up to 90%) of leukemic cells from weeks 8 to 11 post-transplant. Despite the severity of the disease, administration of either VxL chemotherapy or anti-pTα-DM1 ADC resulted in a significant drop of leukemia burden in the PB during the 8-11 weeks treatment period (**Figure7B**). Notably, median survival of treated mice was significantly prolonged when compared to that of control animals, which were euthanized at day 90 (**Figure 7C**). Likewise, tumor progression was markedly impaired in NSG mice treated with the anti-pTα-MMAE ADC, which received T-ALL42 cells with high engraftment potential (disease established by 5 weeks post-transplant, **Figure 7A**). Reduced leukemic cell numbers in treated mice correlated with >60% prolonged median survival, relative to control mice treated with an irrelevant IgG2a-MMAE ADC control (**Figure 7D, E**). Collectively, these results provide evidence for the efficacy of an anti-pTα ADC targeted immunotherapy against pre-TCR+ T-ALL cells with LIC potential, offering a promising strategy to prevent and treat T-ALL relapses.

**Figure 7.**
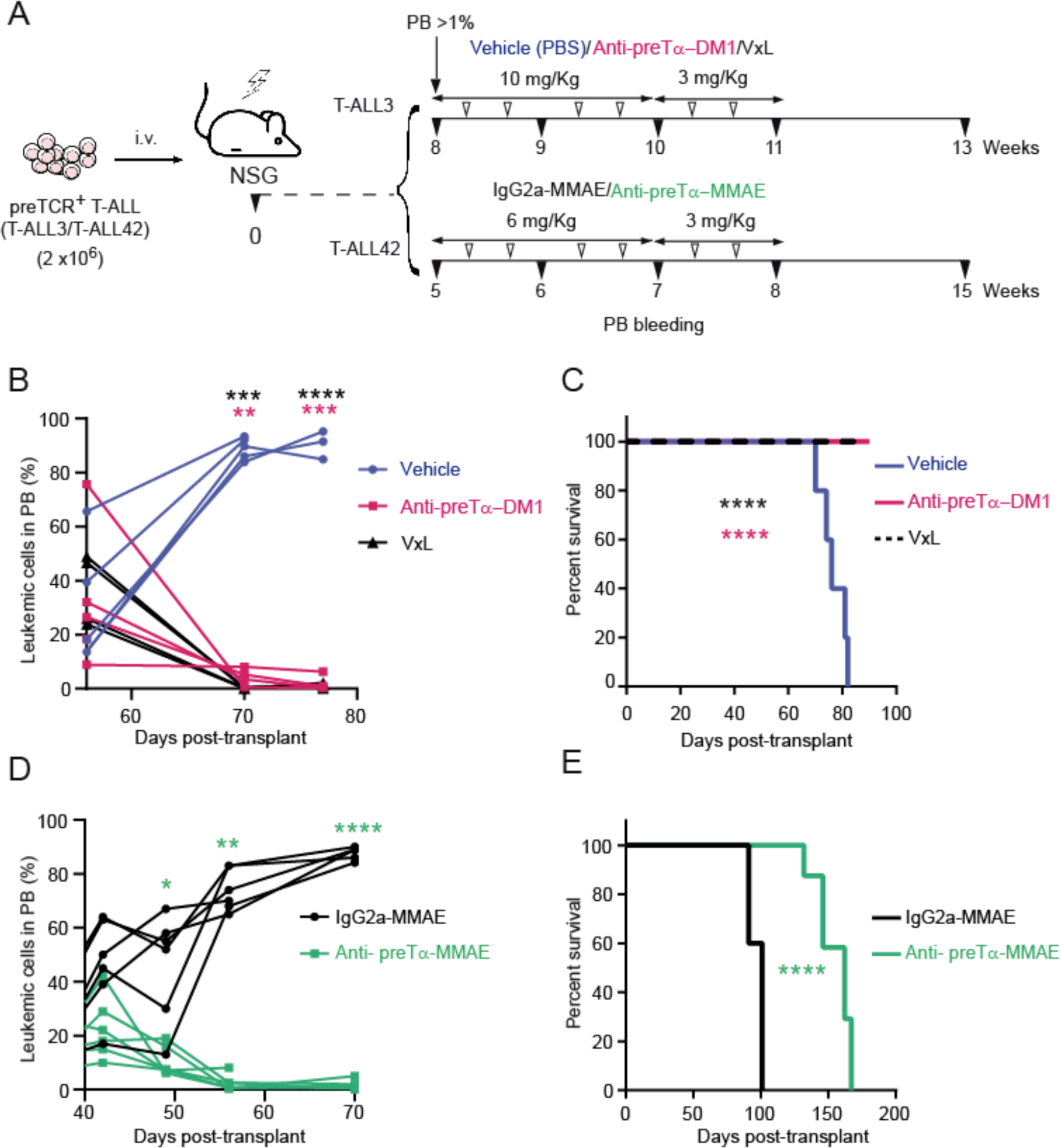
Treatment of patient T-ALL with pre-TCR targeting ADCs impairs tumor progression and increases overall survival in a preclinical *in vivo* assay. (A) Timeline of the *in vivo* pre-TCR targeting immunotherapy based on administration of anti-pTα-DM1 or anti-pTα-MMAE ADCs in NSG mice injected with patient T-ALL3 or T-ALL42. (B, D) Percentages of T-ALL3 (B) or T-ALL42 (D) leukemic cells recovered at the indicated days from the PB of mice treated with either anti-pTα-DM1, vehicle (PBS), or VxL (B), or with anti-pTα-MMAE or IgG2a-MMAE (D). (C, E) Kaplan-Meier survival curves of mice injected with T-ALL3 (C) or T-ALL42 cells (E) and treated as indicated in (B) and (C), respectively. Data are shown as values from 5 (B, C) or 6 (D, E) mice per group. \**P* <0.05; \*\**P*< 0.01 \*\*\**P* < 0.001; \*\*\*\**P* < 0.0001.

## DISCUSSION

In this study, we have identified the pre-TCR as a targetable pathway exhibiting key molecular and functional features optimal for developing selective T-ALL immunotherapy. This finding represents an important advance in the field, as T-ALL targeted therapies have remained underdeveloped owing to challenges in identifying targetable pathways restricted to malignant T cells. Critically, the pre-TCR is not expressed on healthy mature T cells, but it was previously proposed to be present in half of TCRαβ-lineage T-ALL cases that display the phenotype and TCR status of pre-TCR^+^ thymocytes arrested at the β-selection developmental stage (37). Here, we formally demonstrate that patient T-ALLs with a “β-selection” phenotype are reactive with a specific anti-human pTα mAb (39), confirming the existence of ∼50% of human T-ALL cases that express a surface pre-TCR endowed with signaling properties. This discovery provided the molecular basis for developing an anti-pre-TCR ADC targeted therapy, which efficiently impairs T-ALL progression and increases survival in preclinical assays, as proof of concept of a promising immunotherapy for T-ALL patients.

An important aspect uncovered by this study is that pre-TCR expression is a biomarker of T-ALL cells with LIC potential. The therapeutic relevance of this finding is strengthened by the observation that pre-TCR signaling is required for LIC function, a feature that may prevent the loss of cell-surface pre-TCR from T-ALL in response to therapeutic pressure. Pre-TCR-dependency of malignant LIC growth is likely to result from the physiologic proliferative signals mediated by the pre-TCR, rather than from a specific oncogenic effect, suggesting that activation pathways downstream of pre-TCR may be key therapeutic targets. Accordingly, our biochemical and functional data show that several pre-TCR signaling pathways activated at the β-selection stage are functional in T-ALL cells as well (43–46). Although further studies are required to confirm that pre-TCR signaling works similarly in leukemia and developing pre-T cells, it is worth noting that patient T-ALLs recapitulate physiological pre-TCR expression features, with limited numbers of pre-TCR complexes reaching the leukemic cell surface (42). Notably, low pre-TCR expression was sufficient to provide a selective growth advantage to T-ALL cells *in vivo*, a finding with important therapeutic implications regarding the functional efficacy of pre-TCR targeting with specific mAbs. Given that seminal studies recently showed that physiological pre-TCR function involves interaction with endogenous peptides bound to Major Histocompatibility Complex Molecules (pMHC) (51, 52), it is likely that binding to pMHC ligands is also required to induce pre-TCR function and LIC potential of T-ALLs. Therefore, disruption of pre-TCR-pMHC interactions with blocking mAbs may have a therapeutic impact that needs to be directly studied.

Regarding the importance of the pre-TCR at the early induction phase of human T-ALL, it may depend on the particular driving oncogenes, as reported in mice (31–36). Nonetheless, pre-TCR function commonly accelerates leukemia onset and increases leukemia aggressiveness in different T-ALL mouse models (33, 34, 36), a finding which concurs with the observation that human progenitors undergoing pre-TCR-mediated β-selection are particularly prone to malignant transformation (37). Therefore, the pre-TCR may play a cooperative role in distinct human T-ALL oncogenic settings. This is particularly exemplified by oncogenic NOTCH1, as ICN1 constructs with different activation potential can lead to either pre-TCR-dependent (31, **Figure S2**) or -independent (36) T-ALL generation in mice, although pre-TCR signaling largely facilitates NOTCH1-induced leukemogenesis (36), as also does NOTCH3 (32). Given that NOTCH1 activating mutations are found in most molecular T-ALL subgroups (17,18), and that *PTCRA* is a NOTCH1 target (23,24), it is possible that pre-TCR has a common role facilitating leukemogenesis in distinct human T-ALL subtypes with active NOTCH1. Accordingly, pre-TCR expression and NOTCH1 activation are coincident events in our T-ALL patient cohort (**Table S1**).

Overall, the dependency of T-ALL pathophysiology on pre-TCR function provides compelling evidence that the pre-TCR may be an ideal targetable pathway for fighting relapse of distinct T-ALL subtypes. Two additional molecules, CD1a (53) and CCR9 (54) have recently been proposed as suitable therapeutic targets for treatment of (r/r)T-ALL with CAR-T cells. However, no evidence has shown their requirement for T-ALL initiation or progression (53,54), and CD1a was shown to be rarely expressed in (r/r)T-ALL (55). While comprehensive immunophenotypic studies for T-ALL in the relapse setting are needed to confirm pre-TCR expression in (r/r)T-ALL, our preclinical data provide formal proof that pre-TCR targeting by anti-pTα mAbs is a valid therapeutic strategy against LIC cells responsible of T-ALL relapse. In fact, anti-pTα mAb administration led to a marked drop of pre-TCR^+^ T-ALL cells in the PB of SCID mice (not shown), likely involving complement-dependent cytotoxicity (CDC) or Ab-dependent cellular cytotoxicity (ADCC) or phagocytosis (ADCP). In addition, we cannot preclude a direct pro-apoptotic effect of the anti-pTα mAb, as observed *in vitro* (not shown), resulting from persistent pre-TCR engagement and activation. Accordingly, persistent TCR/CD3 signaling of TCRαβ^+^ T-ALLs, experimentally induced by mAbs, has antileukemic properties and leads to massive leukemic cell death in PDX models (56). Whatever mechanism underlies the therapeutic effect of anti-pTα mAb administration, the particular endocytosis and internalization features of the pre-TCR (42,49) suggested that binding of anti-pTα to cytotoxic drugs (ADCs) could provide a more potent therapy. Supporting our hypothesis, we found that anti-pTα binding to DM1 or MMAE, two drugs approved to generate other ADCs currently in the clinic (48), efficiently impaired progression of pre-TCR^+^ T-ALL xenografts, and essentially no T-ALL cells were detectable in the blood of host mice during the treatment period. Therefore, we provided formal proof of the superior efficacy of pre-TCR targeted immunotherapy induced by ADCs as compared with no-drug conjugated antibodies. The safety and efficacy of the ADC strategy for different hematologic tumors have been established in clinical trials (48) and, recently, preclinical data support the ability of anti-TRBC1 ADCs to cure disseminated T-cell cancers (57). Of note, anti-TRBC1–SG3249 was significantly more potent than anti-TRBC1–MMAE treatment (57), suggesting that the SG3249 payload may be also optimal for pre-TCR targeting with ADCs. Other safety aspects to be considered rely on the possible off-target effects of the proposed targeted therapy. Given that the pre-TCR is exclusively expressed within the body in a transient population of thymus-restricted T-cell progenitors (26,30) no severe irreversible toxicities are expected in young infants, in which pre-TCR^-^ progenitors upstream of β-selection could eventually regenerate pre-TCR^+^ thymocytes that would finally mature into functional T cells. Neither severe defects are expected in pediatric and adult patients, which already display a peripheral long-lasting mature T-cell repertoire generated very early in life (58). We thus conclude that the anti-pTα ADC targeting strategy validated in this study holds great promise for therapeutic purposes in patients with (r/r)T-ALL.

## METHODS

### Human and mouse cell samples

Human samples were obtained according to the Declaration of Helsinki principles and to the procedures approved by Local Bioethics Committees after informed consent was provided. Human postnatal thymus samples were removed during corrective cardiac surgery of patients aged 1 month to 4 years. Human fetal thymus samples were obtained from volunteers after legal termination of pregnancy. ETPs were isolated by inmunomagnetic sorting from thymocyte suspensions obtained by centrifugation on Ficoll-Hypaque (Lymphoprep, Axis-Shield PoC AS) using the Dynal CD34 Progenitor Cell Selection System (Life Technologies). Human fetal thymus samples were HPCs were obtained from Ficoll-Hypaque–purified CB samples using the CD34 Progenitor Cell Isolation Kit (Miltenyi Biotec), as described (38). Sorted populations were proved at least 99% CD34^+^ and negative for CD3, CD4, CD8, CD13, CD14, CD19 and CD56 lineage markers on reanalysis (Lin^−^). Primary T-ALL cells were Ficoll-Hypaque–purified from PB or BM patient samples obtained at the time of diagnosis. Mouse HPCs were isolated by cell sorting (FACSAria, BD Biosciences) of cells labelled with APC-coupled anti-CD117 (c-kit) (BD Biosciences), from Lin^-^ BM samples 6- to −10-week-old C57BL/6 mice (The Jackson Laboratory) or C80G mutant mice (47). Lin^−^ cells were obtained by inmunomagnetic sorting using biotin-coupled mAbs against B220/CD45R, Gr1.1, CD11b, CD3, or Ter-119 (BD Biosciences) and streptavidin-coupled magnetic spheres (Miltenyi Biotec), as described (59).

### Flow cytometry

Flow cytometry mAbs included directly-labeled mouse anti-human CD4-PE-Cy5 (clone 13B8.2) and TCR-αβ–PE-Cy5 (clone IP26A) from Beckman Coulter; CD3-PE (clone UCHT1), CD3-APC (clone UCHT1), CD5-FITC (clone L17F12), CD7-FITC (clone 4H9), CD8-PE-Cy7 (clone RPA-T8), CD19-PE (clone HIB19), CD45-FITC (clone HI30), CD45-APC (clone HI30), HLA-DR-PE (clone TV36) and CD7-PE (clone M-T701) from BD Biosciences; CD8-PE (clone 3B5) and TCR-γδ-PE-Cy5 (clone 5A6.E9) from Invitrogen; and CD10-PerCP-Cy5.5 (clone HI10a) from BioLegend. Anti-mouse mAbs included CD8-FITC (clone 5H10) from Invitrogen; CD7-PE (clone 6B7) from Thermo Fisher Scientific; CD44-PE (clone IM7), CD3-PE (clone 145-2C11), CD4-PerCP (clone RM4-5), CD11b-FITC (clone M1/70), Gr1-PE (clone RB6-8C5), H2-Kb-PE (clone AF6-88.5) and H2-Kb-Biotin (clone AF6-88.5) from BD Biosciences; TCRβ-FITC (clone H57-597) from ImmunoTools; CD45R-PE-Cy5 (clone RA3-6B2) from Caltag and CD90 1.2-Pacific blue (clone 30-H12) from Biolegend. For pre-TCR detection, cells were incubated with the affinity-purified anti-pTα mAb K5G3 (39), washed and incubated either with a goat anti-mouse IgG (H+L)-FITC (Invitrogen) or with a biotin-coupled anti-mouse IgG F(abߣ)_2_ (Jackson Immunoresearch). Biotinylated antibodies were developed using PE-Cy7-(Biolegend) or APC-coupled (eBioscience) Streptavidin. Isotype-matched irrelevant Abs (BD Biosciences) were used to define background fluorescence. Flow cytometry was performed in a FACSCanto II (BD Biosciences). Data analysis was performed using FlowJo vX (LLC software).

### Cell lines, OP9-Jag2 cultures and ATOs

Mycoplasma-free JR.pTα and JR.MigR1 T-ALL cell lines generated as described (44) by lentiviral transduction with MigR1-pTα or MigR1-control vectors, respectively, and SupT1, HPB-ALL, Jurkat and CUTLL1 T-ALL cell lines (American Type Culture Collection (ATCC)) were cultured in RPMI 1640 medium (Lonza) supplemented with 10% fetal bovine serum (FBS; Gibco). For *in vitro* T-cell differentiation, HPCs isolated from CB were seeded (10^5^/ml) on semiconfluent monolayers of OP9 stromal cells expressing the human Jag2 Notch ligand (OP9-Jag2) (60) and cultured in p24-well plates with α-MEM medium (Gibco) supplemented with 20% FBS, 200 IU/ml of recombinant human (rh)IL-7, 100 IU/ml of rhSCF (National Institute of Biological Standards and Controls) and 50 ng/ml of rhFLT3L (PeproTech). T-cell development was also analyzed from human ETPs co-cultured with the stromal cell line MS5 transduced with the Notch ligand DLL4 (MS5-hDLL4), in artificial thymic organoids (ATOs) generated as described (61). ATOs were plated in a 0.4 μM Millicell Transwell (Thermo Fisher Scientific) insert and placed in one well of a six-well plate in 1 ml serum free culture medium (“RB27”) composed of RPMI 1640 (Lonza), 4% B27 supplement (Thermo Fisher Scientific), 30 μM L-ascorbic acid 2-phosphate sesquimagnesium salt hydrate (Sigma-Aldrich), 1% Glutamax (Thermo Fisher Scientific), 200 IU/ml of rhIL-7 and 5 ng/ml of rhFlt3-L.

### T-ALL generation, xenotransplantion, and LIC activity assay

All animal studies were approved by the Animal Experimentation Ethics Committee of the Comunidad de Madrid Animal Research Review Board (PROEX002621). Human T-ALL was generated as described (38) in sublethally irradiated (1.5 Gy) 6- to 10-weeks-old NOD.Cg-Prkdcscid Il2rgtm1Wjl/SzJ (NSG; The Jackson Laboratory) mice subjected to i.v. injection with human CB HPCs transduced with the ICN1 construct (62). Mouse T-ALL was generated as reported (59), in NSG mice subjected to i.v. injection with mouse (C57Bl/6) BM Lin-c-kit+ HPCs transduced with ICN1, following a modification of the original model (40). Human *in vivo* T-cell development was analyzed in sublethally irradiated (3,5 Gy) 3- to 5-days-old RAG-2^-/-^ x γc ^-/-^ mice (63) subjected to intrahepatic injection with shCMS- or shSC-transduced human ETPs (10^5^). At 3- and 5-weeks post-transplant, thymuses from host mice were collected and analyzed for reconstitution with human developing T cells by flow cytometry. For leukemia xenotransplantation assays, sublethally irradiated (1.5 Gy) 6- to 10- week-old NSG mice were subjected to i.v. injection with T-ALL or B-ALL patient samples or with human T-ALLs generated de novo in host mice (first grafts). When indicated, first grafts from host mice were i.v. injected into secondary recipients (second grafts). LIC activity was analyzed by transplantation into NSG mice under limiting dilution conditions (10^5^ to 10 cells per mouse, 1-4 mice per dose) as reported (59). Transplanted mice were analyzed by flow cytometry for the presence of human CD7^+^ CD45^+^ blasts in PB from 6 to 32 weeks post-transplant, and the frequency of leukemia-free mice was calculated using the ELDA software (http://bioinf.wehi.edu.au/software/elda/) (64).

### shRNA lentiviral constructs and transductions

For silencing endogenous CMS expression, 5 distinct shRNAs directed against *CD2AP* encoding human CMS (shCMS) (Sigma-Aldrich Mission TRCN shRNA Target set, TRCN0000119097, TRCN0000119098, TRCN0000119099, TRCN0000119100, TRCN0000119101) were assayed by transfection and puromycin selection (1 μg/ml; Sigma-Aldrich) of SupT1 cells, then analyzed for CMS expression by western blotting. The best-performing shCMS or a scrambled shRNA (shSC), used as control, were then cloned under the U6 promoter into the pHRSIN-GFP lentiviral vector. T-ALL and B-ALL cell transduction was performed as described (59) in the presence of rhIL-7 (200 IU/mL).

### Western blotting

Activation of signaling pathways downstream of pre-TCR was analyzed by western blotting of cells incubated with 10 µg/ml of anti-CD3ε (UCHT-1; BD Biosciences) or anti-pTα (K5G3) (39) mAbs at 4°C for 30 min, followed by crosslinking with 10 µg/ml of anti-mouse IgG (Sigma-Aldrich) at 37°C for the indicated incubation times. Whole-cell lysates (RIPA buffer) separated on 10% sodium dodecyl sulfate-polyacrylamide (Bio-Rad) gel electrophoresis (SDS-PAGE) were transferred to polyvinylidene difluoride membranes (Merck Life Science) that were incubated with Abs against either phospho 44/42 MAPK (ERK1/2), p44/42 MAPK ERK (Cell signaling) or CMS (Santa Cruz Biotechnology). α-tubulin expression (Sigma-Aldrich) was analyzed as loading control. For PLCγ activation analysis, cells were lysed in 1 ml Brij96 lysis buffer containing protease and phosphatase inhibitors, and subjected to immunoprecipitation with anti-PLCγ mAb (BD Bioscience) and protein G-sepharose CL-4B beads (Sigma-Aldrich). After SDS-PAGE and immunoblotting, membranes were probed with anti-phospho-Y783-PLCγ Ab or anti-PLCγ mAb (Cell Signaling). Probed membranes were washed and incubated with horseradish peroxidase-conjugated anti-mouse or anti-rabbit Abs for 1 hour and developed with Lumi-LightPLUS Western Blotting Substrate (Roche). Quantification was performed on Lumi-Light autoradiography films using ImageJ software.

### NFAT transcriptional activation assay

For NFAT activation assays, JR.pTα cells (44) were transfected with a luciferase reporter plasmid containing 3 tandem copies of the composite NFAT/AP1-response element from the human *IL-2* gene promoter, together with a pRL-CMV *Renilla* reporter construct (Promega). After 24 hours, cells were cultured for 6 hours with or without plate-bound anti-CD3ε (OKT3, Sigma Aldrich) and lysed using the Dual luciferase assay Kit (Promega). Luciferase activity was normalized to the Renilla luciferase activity and expressed as fold-induction relative to the basal activity of non-stimulated cells, as described (44).

### Anti-pTα mAb and ADC *in vivo* administration

For *in vivo* administration of anti-pTα, CB17/lcr-*Prkd^scid^*/lcrlcoCrl (SCID) immunodeficient mice xenotransplanted with a patient T-ALL (10^6^ T-ALL3 cells per mice) were serially i.p. injected (twice weekly) for 10 weeks with 200 μl of PBS containing either anti-anti-pTα mAb K5G3 (39) or IgG2a isotypic control (10mg/Kg) starting when T-ALL engraftment in PB was >1% (week 3). Anti-pTα ADCs were generated by the Custom Antibody Service from Institute for Advanced Chemistry of Catalonia. Anti-pTα mAb K5G3 (39) was conjugated to Mertansine toxin (DM1) via a non-cleavable MCC linker, and to Monomethyl Auristatin E (MMAE) via a valine-citruline protease-cleavable linker (SC-VC-PABC). The bioconjugation process involves the binding of the drug-linker complex to the lysines of the antibody through the succinimidyl group incorporated at the beginning of both linkers. Briefly, after stirring for 2h at room temperature plus overnight at 4 °C, the resulting mixture was desalted using a compact liquid chromatography system to purify the bioconjugate by using 10 mM phosphate-buffered saline as elution buffer (10 mM sodium phosphate, 137 mM sodium chloride, 3.2 mM potassium chloride, 2 mM potassium dihydrogen phosphate, pH 7.4). For anti-pTα-DM1 treatment, xenotransplanted NSG mice (2×10^6^ T-ALL3 cells) were i.p. injected with either anti-pTα-DM1 (10 mg/Kg, twice weekly, for 2 weeks, and 3 mg/Kg for an additional week) or with vehicle (PBS), starting when T-ALL engraftment in PB was >1% (week 8). A group of mice received a standard VxL induction treatment consisting of vincristine (Selleckchem; 0.15 mg/kg, weekly for 3 weeks), plus dexamethasome (Sigma-Aldrich; 5 mg/Kg) and L-asparaginase (Kidrolase; EuSA Pharma; 1000 U/Kg, weakly during 5 days, for 3 weeks. For anti-pTα-MMAE treatment, xenotransplanted NSG mice (2×10^6^ T-ALL42 cells) were i.p. injected with either anti-pTα-MMAE or IgG2a-MMAE (6 mg/Kg, twice weekly, for 2 weeks, and 3 mg/Kg for an additional week), starting when T-ALL engraftment in PB was >1% (week 5).

### Pre-TCR labeling and internalization of pTα-ADC

SupT1 cells were incubated with 1 µg of anti-pTα mAb (39) or anti-pTα ADC (anti-pTα-DM1 or anti-pTα-MMAE) at 4°C for 30 min, then washed with ice-cold PBS to remove unbound antibody, and incubated at 37°C for the indicated times. Then, cells were washed with ice-cold PBS and the primary antibody was detected using FITC-conjugated goat anti-mouse IgG (H+L) antibody (Invitrogen). Relative pre-TCR surface pre-TCR expression at each time point was calculated as the percentage of positive cells normalized to time 0.

### Cell proliferation and cell cycle analysis

Cells were seeded at 2×10^5^ cells/ml per well in 48 well-plates in the absence or presence of 5 µg/ml of either anti-pTα mAb (39), anti-pTα-DM1, anti-pTα-MMAE, or isotype-matched controls at 37°C in 5% CO_2_. After 4 days, cells were collected, washed and counted. Cell proliferation was calculated as percentages of total cells at the end of culture relative to input cell number. For cell cycle analysis, 4×10^5^ cells were incubated with 5 µg/ml of Hoechst 33342 (Sigma) at 37°C for 45 min. Then, cells were washed with ice-cold PBS and analyzed by flow cytometry.

### Statistical analysis

Statistical significance was determined using a 2-tailed Student t test, and Kaplan-Meyer survival curves were compared using the log-rank test, using GraphPad software. In all cases, significance was defined as p < 0.05.

## ACKNOWLEDGEMENTS

The authors thank J. C. Aster, K. Weijer, H. Spits, Gay Crooks and J. C. Zúñiga-Pflücker, for providing useful reagents and mouse strains; Pablo Menéndez and Clara Bueno (Josep Carreras Leukemia Research Institute) for fetal thymus samples and Eloísa Castillo for technical support. We thank the expert personnel within the Animal Facility and the Flow Cytometry Core at the Centro de Biología Molecular Severo Ochoa and within the Mouse Embryo Cryopreservation Facility at CNB-CSIC for their invaluable assistance, and the Centro de Transfusión de la Comunidad de Madrid, the Pediatric Cardiosurgery Units from Ciudad Sanitaria La Paz and Hospital Universitario 12 de Octubre (Madrid, Spain) for cord blood and postnatal thymus samples, respectively. This research was supported in part by grants PID2019-105623RB-I00, PID2022-138880OB-I00 and PDC2021-121238-I00 funded by Ministerio de Ciencia, Innovación y Universidades. Agencia Estatal de Investigación (MCIN/AEI/ 10.13039/501100011033)/ European Regional Development Fund, European Union, and by grants from Fundación Unoentrecienmil and Fundación Inocente Inocente. Institutional grants from the Fundación Ramón Areces and Banco de Santander to the Centro de Biología Molecular Severo Ochoa are also acknowledged.

## CONTRIBUTIONS

M.L.T. and P.F. conceived and designed the project, supervised experiments and wrote the paper. P.F., M.G.P., M.M., C.C., J.A. and B.A. performed and analyzed experiments; M.C. performed clinical evaluations of patients and provided patient samples.

## Notes

### Competing Interest Statement

The authors have declared no competing interest.

