## Supplementary material for "Pre-TCR-Targeted Immunotherapy for T-cell Acute Lymphoblastic Leukemia": Suplemental Fig1, Fig2 and Table 1

### SUPPLEMENTAL DATA

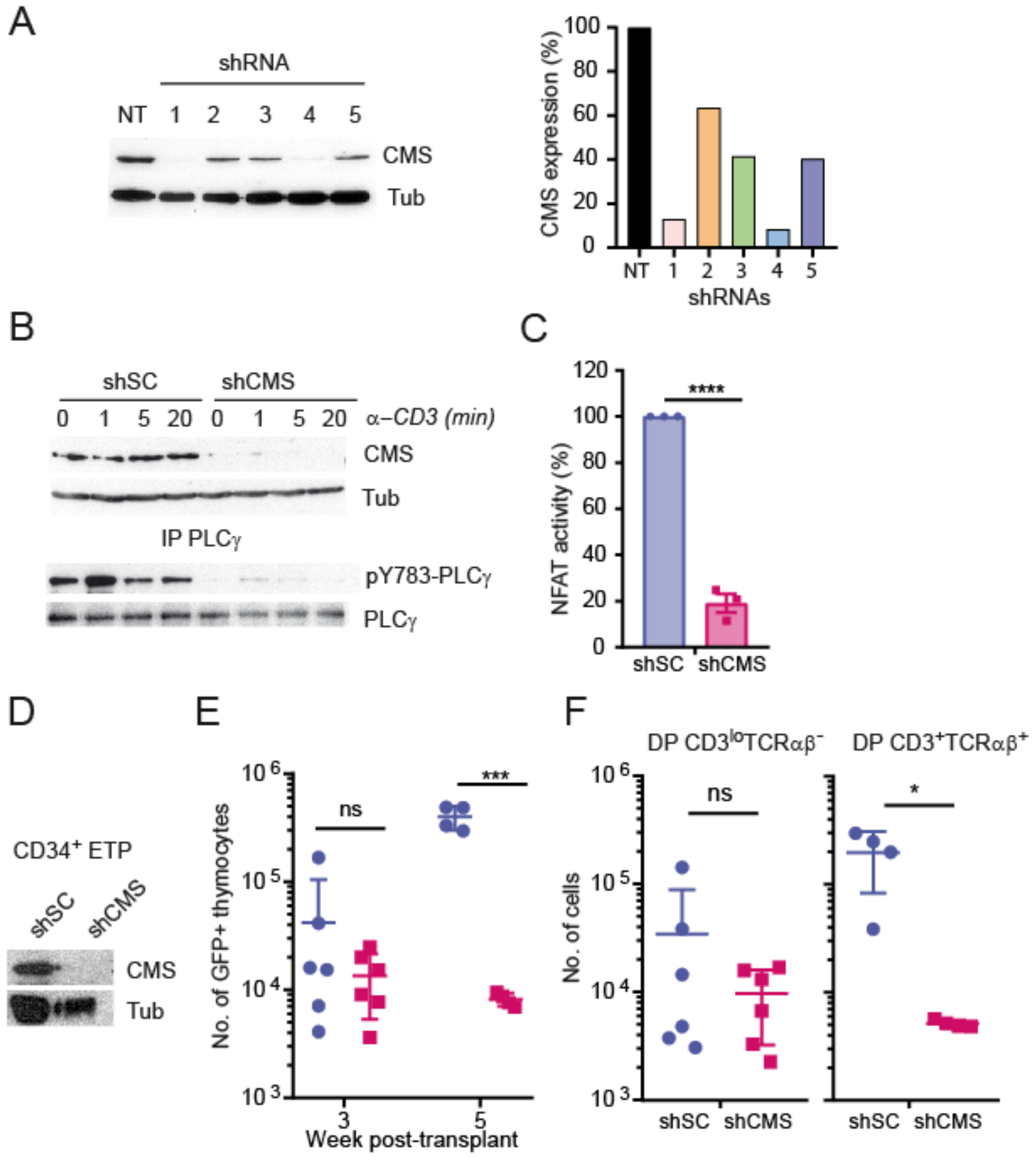


**Figure S1: *In vitro* and *in vivo* abrogation of pre-TCR function by gene silencing of the CMS (*CD2AP*) functional adaptor of pTα.** (A) Immunoblot analysis of CMS expression in SupT1 cells that were either transfected with several shRNAs (shCMS1-5) specific for *CD2AP,* the gene encoding CMS, or left untransfected (NT) (left panel). Relative CMS expression normalized to α-tubulin expression used as loading control is shown in the right panel. (B) Immunoblot analysis of CMS expression (upper) and PLCγ tyrosine (Tyr) phosphorylation (bottom) in JR.pTα pre-TCR$^+$ cells transduced either with shCMS4 or with shSC as control, prior to activation with an anti-CD3ε mAb for the indicated times. PLCγ Tyr-phosphorylation was analyzed after immunoprecipitation with anti-PLCγ by probing with an anti-Y783-PLCγ antibody. Tubulin expression was analyzed as loading control. (C) Relative NFAT activity of JR.pTα cells transducted with either shCMS4 or shSC as control, analyzed upon stimulation with an anti-CD3ε mAb. Data are shown as mean percentages ± SEM of NFAT activity relative to activated shSC cells (n=3). (D) Immunoblot analysis of CMS in human CD34$^+$ early thymic progenitors (ETPs) transducted with a lentivirus encoding either shCMS4 and GFP or shSC and GFP as control (E) Absolute numbers of shCMS- or shSC-transduced ETP-derived human cells reconstituting the thymus of RAG-2$^{-/-}$ x γc$^{-/-}$ mice at the indicated weeks post-transplant. ETP transduction efficiencies with shCMS and shSC were 24,4% ± 4,1% and 21,6% ± 3,5%, respectively. (F) Absolute numbers of shCMS- or shSC-transduced human thymocytes in (E) expressing the DP CD3$^{lo}$ TCRαβ$^-$ phenotype at 3 weeks post-transplant (left panel) or the post-β selected DP CD3$^+$ TCRαβ$^+$ phenotype at 5 weeks post-transplant (right panel). Data in (E, F) are shown as mean numbers ± SEM of transduced (GFP+) cells normalized to $10^5$ transduced input cells intrahepatically injected into 4-6 mice per group in two independent experiments. *$P$ <0.05; ****$P$ < 0.0001.

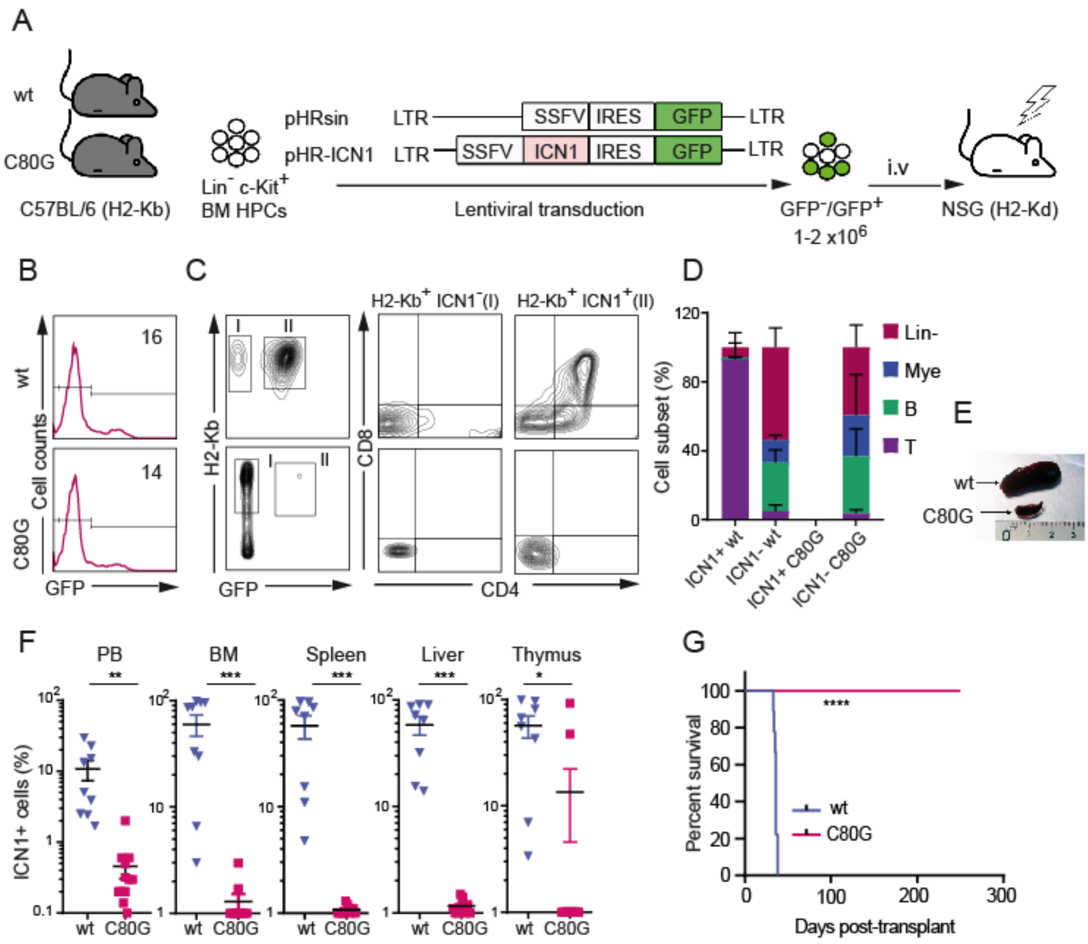


**Figure S2**

(A) Schematic diagram of experimental design for mouse T-ALL generation. Sublethally irradiated NSG (2H-Kd⁺) mice were subjected to i.v. injection with BM lin⁻ c-kit⁺ HPCs isolated from wild type (wt) or CD3ε C80G mutant C57BL/6 (H2-Kb⁺) mice and transduced with a lentiviral vector encoding either active NOTCH1 (ICN1) and GFP or GFP alone as control. (B) Representative flow cytometry analysis of transduction efficiencies of wt (upper) and mutant (bottom) HPCs in (A). Numbers indicate percentages of transduced (GFP⁺) cells. (C) Representative CD8 and CD4 expression of electronically-gated mock-transduced (gate I; ICN1⁻) and ICN1-transduced (gate II; ICN1⁺) cells, derived from donor wt (upper) or C80G mutant (bottom) BM HPCs engrafting the BM of NSG mice at 5 weeks post-transplant. Results are representative of one out of at least 9 mice per group (n=2). (D) Mean percentages ± SEM of human myeloid-lineage ($CD11b^+$ or $Gr1.1^+$), B-lineage ($B220^+$), T-lineage (DP; $CD4^+$ $CD8^-$ and/or $CD8^+$ $CD4^-$) and $Lin^-$ ($CD11b^-$, $Gr1.1^-$, $B220^-$, $CD4^-$, $CD8^-$) cells derived from $ICN1^+$ and $ICN1^-$ wt or C80G mutant BM HPCs engrafting the BM of NSG mice in (C). (D) Image of representative spleens obtained from mice shown in (C, D). (F) Mean percentages ± SEM of $ICN1^+$ cells derived from wt or C80G mutant HPCs engrafting the indicated organs of NSG mice (9 mice per group) at sacrifice. (G) Kaplan-Meier survival curves of transplanted NSG mice in (F). $^{**}P < 0.01$; $^{***}P < 0.001$; $^{****}P < 0.0001$.

**Table S1. Phenotype of human T-ALLsamples**

| Name | EGIL classification | pre-TCR | ICN1 |
|---|---|---|---|
| T-ALL1 | Mature | - | - |
| T-ALL2 | Cortical | - | + |
| T-ALL3 | Pre-T | + | + |
| T-ALL4 | Cortical | + | nd |
| T-ALL5 | Cortical | + | + |
| T-ALL8 | Mature | - | + |
| T-ALL9 | Cortical | + | + |
| T-ALL10 | Pre-T | - | + |
| T-ALL17 | Pre-T | + | + |
| T-ALL18 | Cortical | + | + |
| T-ALL26 | Pre-T | - | nd |
| T-ALL32 | Mature | - | nd |
| T-ALL36 | Cortical | - | nd |
| T-ALL37 | Pre-T | - | nd |
| T-ALL39 | Cortical | + | nd |
| T-ALL40 | Cortical | - | nd |
| T-ALL41 | Cortical | - | nd |
| T-ALL42 | Cortical | + | nd |
| T-ALL43 | Cortical | + | nd |

nd: not determined
